# Transforming growth factor-β promotes the postselection thymic development and peripheral function of interferon-γ-producing invariant natural killer T cells

**DOI:** 10.1101/2022.10.23.513409

**Authors:** Roxroy C. Morgan, Cameron Frank, Munmun Greger, Mikael Sigvardsson, Elizabeth T. Bartom, Barbara L. Kee

## Abstract

Interferon-γ producing invariant natural killer T (iNKT1) cells are lipid reactive innate-like lymphocytes that are resident in the thymus and peripheral tissues where they protect against pathogenic infection. The thymic functions of iNKT1 cells are not fully elucidated but subsets of thymic iNKT cells modulate CD8 T cell, dendritic cell, B cell and thymic epithelial cell numbers or function. Here we show that a subset of thymic iNKT1 cells require transforming growth factor (TGF)-β induced signals for their development and for expression of residency associated adhesion receptors. Liver and spleen iNKT1 cells do not share this TGF-β gene signature but nonetheless TGF-β is required for optimal liver iNKT1 cell function. Our findings provide insight into the heterogeneity of mechanisms guiding iNKT1 cell development in different tissues and suggest a close association between a subset of iNKT1 cells and TGF-β producing cells in the thymus.

## Introduction

Invariant natural killer T (iNKT) cells are a subset of innate like T lymphocytes that express an invariant T cell receptor that recognizes glycolipid antigens in the context of the non-classical major histocompatibility complex protein CD1d (Shissler and Webb, 2019). These cells comprise a large portion of T lymphocytes in peripheral tissues. As much as 40% and 5% of liver T cells are iNKT cells in mice and humans respectively (Crosby and Kronenberg, 2018; Syn et al., 2010). The majority of liver iNKT cells are tissue resident, interferon (IFN)-γ producing iNKT1 cells (Crosby and Kronenberg, 2018; Lee et al., 2015). These cells patrol the sinusoidal space to protect against invading pathogens such as the spirochete *Borrelia burgdorferi*, the causative agent of Lyme’s Disease, and viral infection such as Hepatitis C Virus. (Crosby and Kronenberg, 2018; Thomas et al., 2011; Umeshappa et al., 2022). Chronic activation of iNKT cells is associated with disease severity in nonalcoholic fatty liver disease (NAFLD) and nonalcoholic steatohepatitis (NASH) (Crosby and Kronenberg, 2018; Syn et al., 2012; Syn et al., 2010). iNKT cells have also been implicated in protection from pathogens in the lung and other tissues but may contribute to inflammatory disease such as asthma and atherosclerosis.

iNKT cell development initiates in the thymus from CD4^+^CD8^+^ thymocytes after rearrangement of the iNKT cell receptor (Bendelac et al., 2007; Benlagha et al., 2002). During selection, iNKT cells are recruited to the thymic medulla by CCL21 and a subset of these immature postselection iNKT cells emigrate to seed peripheral tissues where they continue their maturation (Baranek et al., 2020; Harsha Krovi et al., 2020; Wang and Hogquist, 2018). The cells remaining in the thymus differentiate into multipe effector subsets but the majority are thymic resident IFN-γ-producing iNKT1 cells (Engel et al., 2016; Lee et al., 2016). Cytokines produced by thymic iNKT cells, in particular IL-4, have the potential to impact the function of multiple cell types including CD8 T cells and dendritic cells (Breed et al., 2022; Verykokakis et al., 2010; Weinreich et al., 2010). While our understanding of of iNKT cell selection is well developed, the mechanisms that promote postselection iNKT1 cell differentiation in different tissues is not well understood.

Here, using a novel model for Cre-mediated recombination in postselection iNKT1 cells, we demonstrate that thymic iNKT1 cells differ from liver and spleen iNKT1 cells by expression of genes associated with transforming growth factor (TGF)-β signaling and reduced expression of genes associated with cytokine signaling. While TGF-β is known to regulate iNKT cell selection (Doisne et al., 2009), we demonstrated that TGF-β signaling in postselection thymic iNKT1 cells enforced a classic TGF-β-associated gene signature, including repression of migratory genes and induction of multiple residency-associated adhesion receptors and repressed a cytokine signaling gene signature. Postselection iNKT1 cells that cannot respond to TGF-β due to deletion of *Tgfbr2* remained resident but failed to differentiate into cells that expressed the residency-associated adhesion receptors CD49a and CD103. In contrast to studies in which *Tgfbr2* was deleted in other IFN-γ producing cell types, including preselection iNKT cells (Doisne et al., 2009), NK cells (Viel et al., 2016), or CD8 T cells differentiated into tissue resident memory cells (Trm) (Crowl et al., 2022; Mackay et al., 2013; Mackay et al., 2015), we found no role for TGF-β signaling in the control of the transcription factor T-BET or T-BET target genes. Despite the lack of a TGF-β gene signature in liver iNKT1 cells, TGF-β signaling was required for their optimal expression of CD49a and supported their production of IFN-γ and IL-4. Our data reveal a selective requirement for TGF-β signaling in the postselection generation of thymic CD49a^+^CD103^+^ iNKT1 cells and optimal liver iNKT1 function.

## Results

### Thymic iNKT1 cells express genes associated with TGF-β signaling

To trace postselection iNKT1 cells in different tissues we crossed a *Tbx21^Cre^* bacterial artificial chromosome transgenic mouse to Rosa26-Stop-floxed-YFP mice (Haddad et al., 2013; Srinivas et al., 2001). *Tbx21* encodes T-BET, the signature transcription factor for iNKT1 cell, and therefore we anticipated that iNKT1 cells and possibly their effector fate restricted progenitors would be YFP^+^. Indeed, we found that YFP^+^ cells in the thymus, liver and spleen were primarily CD44^+^NK1.1^+^, a phenotype associated with iNKT1 cells (Fig. S1A). We used this model to isolate iNKT1 cells from the thymus, liver, and spleen and compare their gene programs by RNA-sequencing. A prior study demonstrated that iNKT1 cells isolated from different tissues, using a different selection strategy, had essentially identical gene programs (Murray et al., 2021). Consistent with this, we found that the transcriptome of liver and spleen iNKT1 cells were nearly identical with < 20 differentially expressed genes (DEGs) (Fig. 1A). However, the transcriptome of thymus and liver iNKT1 cells differed by 124 genes (adj.p < 0.01) whereas thymic and splenic cells differed by 107 genes (Fig. 1A). While these are small differences, there were specific genes of interest among the DEGs. Using Gene Set Enrichment Analysis (GSEA), we observed a significant enrichment (FDR <25%) of multiple Hallmark pathways in the liver as compared to the thymus including in IL6-JAK-STAT3 signaling, mTORC1 signaling and fatty acid metabolism (Fig. 1B, S1B). As expected, similar GSEA pathways were enriched in the liver and the spleen when compared to the thymus (Fig. S1B).

**Figure 1:**
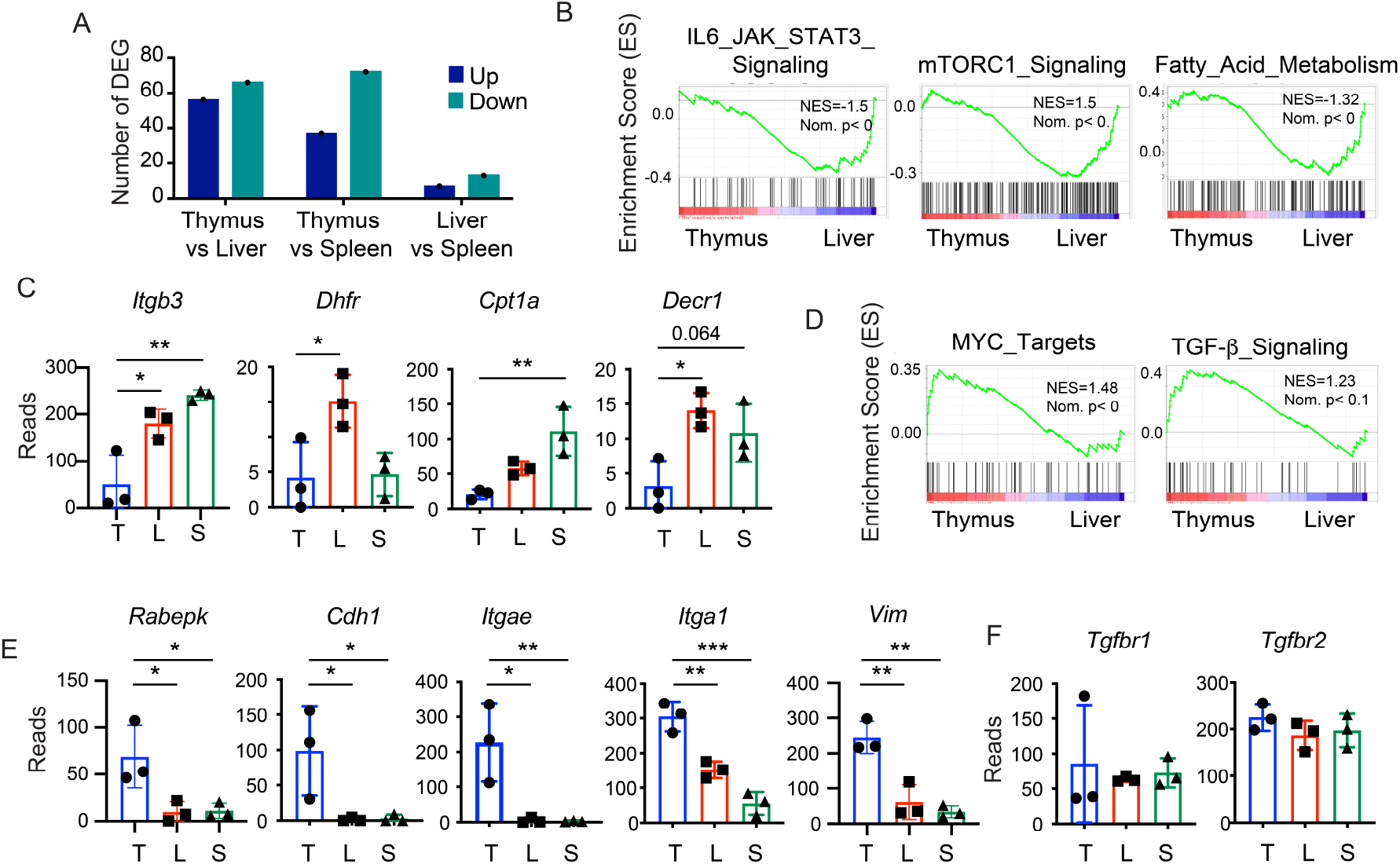
Thymic iNKT1 cells are enriched for genes associated TGF-β signaling compared to liver and spleen iNKT1 cells. RNA was sequenced from WT iNKT1 cells isolated from the thymus, liver and spleen and normalized read counts were compared. (A) Number of differentially expressed genes (DEG) between WT thymus, liver and spleen iNKT1 cells. (B) Hallmark Pathways identified by GSEA as enriched in liver compared to thymus iNKT1 cells. (C) Reads for 3 replicate RNA-seq samples from WT thymus (T, blue), liver (L, red) and spleen (S, green) for *Itgb3, Dhfr, Cpt1a* and *Decr*, which are representative genes from the pathways identified in (B). (D) Hallmark Pathways identified as enriched in thymus as compared to liver iNKT1 cells. (E) Normalized reads from replicate RNA-seq samples showing genes representative the pathways identified in (D). (F) RNA-seq reads for *Tgfbr1* and *Tgfbr2.* *P<0.05, **<P0.01, ***P<0.005 by ANOVA with multiple comparisons.

In the IL6-JAK-STAT3 pathway, *Itgb3* was higher in liver and spleen compared to the thymus iNKT1 cells whereas *Socs3* mRNA, encoding a suppressor of cytokine signaling that binds STAT3, did not quite reach significance (Fig. 1C, S1C). In the mTORC1 pathway, which was not enriched in the thymus to spleen comparison, *Dhfr* mRNA was highest in the liver (Fig. 1C, S1C). Fatty acid metabolism is required for survival of CD8 Trm cells in the skin (Pan et al., 2017), and CD8 Trm cells in different tissues utilize distinct fatty acid binding proteins to uptake fatty acids (Frizzell et al., 2020). We observed that mRNA encoding enzymes in this pathway, including *Cpt1a* and *Decr1* mRNAs, encoding carnitine palmityoyltransferase 1A and 2,4 dienoyl-CoA reductase respectively, were higher in both the liver and spleen compared to the thymus even though GSEA identified this pathway as enriched only in the liver (Fig. 1C, S1C). These data suggest that liver and spleen iNKT1 cells have active IL6-JAK-STAT3 signaling and may utilize fatty acid metabolism but that these pathways are less active in thymic iNKT1 cells. Interestingly, thymic iNKT1 cells were significantly enriched for only the hallmark MYC-Targets pathway but only *Rabepk*, encoding a mannose 6-phosphage receptor transport protein, reach statistical significance (Fig. 1D, E). In contrast, the TGF-β signaling pathway appeared tobe enriched in thymic iNKT1 cells did reach statistical significance by GSEA; however, multiple genes in this pathway were differentially expressed by RNA-seq (Fig. 1D, E). These genes include *Cdh1*, encoding Cadherin, *Itgae,* encoding the alpha chain of the integrin CD103, *Itga1*, encoding the alpha chain of CD49a, and *Vim*, encoding vimentin (Fig. 1D). *Tgfbr1* and *Tgfbr2*, encoding components of the TGF-β receptor, were expressed similarly in all iNKT1 cells suggesting that differences in TGF-β signaling-associated genes are not a consequence of TGF-β receptor expression (Fig. 1F). *Tbx21*, encoding the transcription factor T-BET, which is required for development of iNKT1 cells and CD8 Trm cells but is dampened by TGF-β signaling in many cell types, was also not differentially expressed (Fig. S1C) (Crowl et al., 2022; Li et al., 2006b; Mackay et al., 2015; Nath et al., 2019; Townsend et al., 2004). These data indicate that the transcriptome of iNKT1 cells is very similar in the thymus, liver and spleen, but there are DEGs associated with cytokine signaling, metabolism, and TGF-β signaling.

### Differential expression of adhesion receptors on thymic and liver iNKT1 cells

The enrichment of TGF-β signaling induced transcripts was of interest given that iNKT1 cells are tissue resident cells with hybrid NK cell, ILC1, and CD4 and CD8 Trm cell characteristics (Verykokakis et al., 2014). Response to TGF-β depends on *Runx3*, which is expressed in CD8 Trm but not CD4 Trm, resulting in a differential response to this cytokine (Fonseca et al., 2022). Our RNA-seq data revealed a modest increase in *Runx1* mRNA in liver and spleen as compared to thymus iNKT1 cells and broad expression of *Runx3* mRNA in all iNKT1 cells, consistent with their ability to respond to TGF-β (Fig. S1C). *Itgae* and *Itga1*, components of CD103 and CD49a respectively, are induced on subsets of CD8 Trm cells by TGF-β produced from epithelial cells (Fonseca et al., 2022; Hadley et al., 1999; Mokrani et al., 2014; Zhang and Bevan, 2013). By flow cytometry we confirmed that CD103 was expressed on a subset of iNKT1 cells in the thymus but not in the liver (Fig. 2A, B). By contrast, CD69, a protein expressed on tissue resident cells that functions as an inhibitor of the S1P receptor S1P1 (Shiow et al., 2006; Stein et al., 2021), was highly expressed on both thymic and liver iNKT1 cells with a subtle increase on liver iNKT1 cells (Fig. 2A, B). CD49a was expressed on both thymic and liver iNKT1 cells although at a lower frequency and lower MFI on liver iNKT1 cells (Fig. 2A, C). A substantial population of thymic iNKT1 cells expressed CD49a along with CD103 and CD69 (Fig. 2A). These receptors are associated with tissue residency of CD8 T cells; however, liver iNKT1 cells are known to require LFA-1 and ICAM-1 to maintain tissue residency (Thomas et al., 2011). Notably, the MFI for ICAM-1 and LFA-1 on liver iNKT1 cells was higher than on thymic iNKT1 cells (Fig. 2D, E). CD44, another adhesion receptor associated with mature iNKT cells (Benlagha et al., 2005), was also expressed at slightly higher levels on liver as compared to thymic iNKT1 cells (Fig. 2F). These data reveal striking differences in the tissue adhesion programs of thymic and liver iNKT1 cells and are consistent with a TGF-β induced program in the thymus.

**Figure 2:**
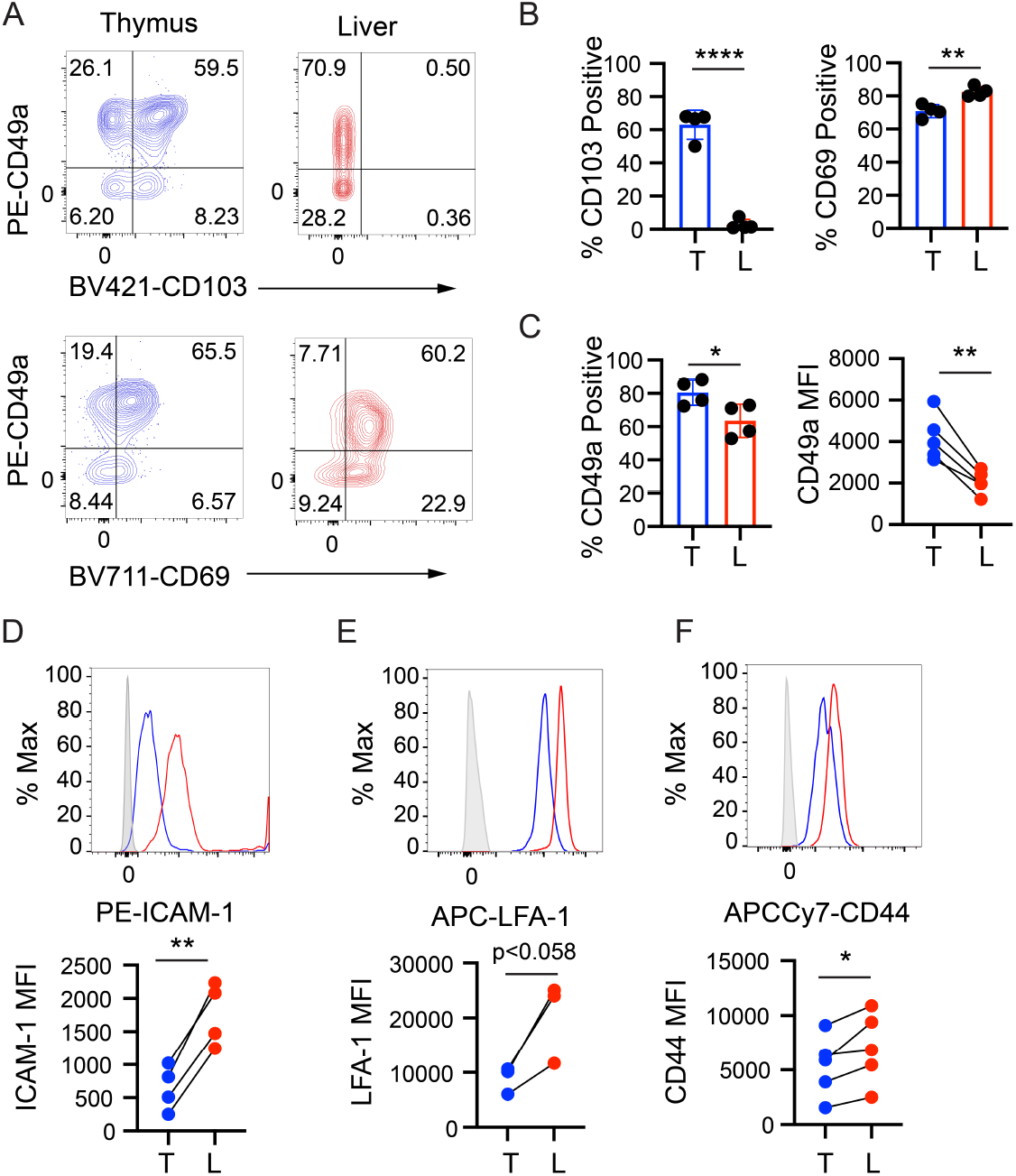
Thymic iNKT1 cells uniquely express TGF-β-associated adhesion proteins. (A) Flow cytometry for CD49a versus CD103 (top) or CD69 (bottom) on iNKT1 cells from the thymus (blue) or liver (red). n = 4 (B) Summary of multiple flow cytometry experiments showing the percent of iNKT1 cells that are positive for CD103 or CD69 or (D) CD49a in the thymus (blue) or liver (red). MFI for CD49a is shown in the right panel. Each data point is an independent mouse. Representative flow cytometry histograms (top) and summary of multiple experiments (bottom) for (D) ICAM-1, (E) LFA-1 or (F) CD44 on thymus (blue) or liver (red) iNKT1 cells. Data are representative of 3 experiments. *P<0.05, **<P0.01, ***P<0.005 by Student’s t-test.

### TGF-β signaling is required for development of CD49a^+^CD103^+^ thymic iNKT1 cells

Previous studies demonstrated a requirement for TGF-β during selection of iNKT cells and a possible role for TGF-β in promoting iNKT17 differentiation while limiting iNKT1 differentiation (Doisne et al., 2009; Havenar-Daughton et al., 2012; Li et al., 2006a). However, in those studies, *Tgfbr2* was inactivated in all T cells prior to T cell receptor-mediated selection making it difficult to dissociate effects on selection from effects on differentiation. To address the role of TGFβRII on iNKT1 cell development postselection, we crossed *Tbx21^Cre^ Rosa26-StopFlox-YFP* mice to mice with floxed alleles of *Tgfbr2*, encoding TGFβRII (Srinivas et al., 2001), to create *Tgfbr2^F/F^ Tbx21^Cre^ Rosa26-StopFlox-YFP* (cKO) mice. In these mice, the total number of YFP^+^ iNKT cells (iNKT1) in the thymus was decreased by 30% compared to *Tbx21^Cre^ Rosa26-StopFlox-YFP* (Ctrl) mice (Fig. 3A. B). The iNKT1 cells remaining in the cKO thymus lacked expression of CD103 but the frequency of CD49a^+^CD103^-^ cells was not increased suggesting that the development of CD49a^+^CD103^+^ iNKT1 depended on TGFβ signaling (Fig. 3C, D). The MFI of CD49a on CD49a^+^CD103^-^ iNKT1 cells was reduced indicating that TGFβRII was also required for proper expression of CD49a (Fig. 3C, D). Interestingly, CD69 was expressed on the majority of cKO iNKT1 cells but the frequency of CD49a^-^CD69^-^ cells was increased, consistent with the loss of some CD49a^+^CD69^+^iNKT1 cells (Fig. 3C, D).

**Figure 3:**
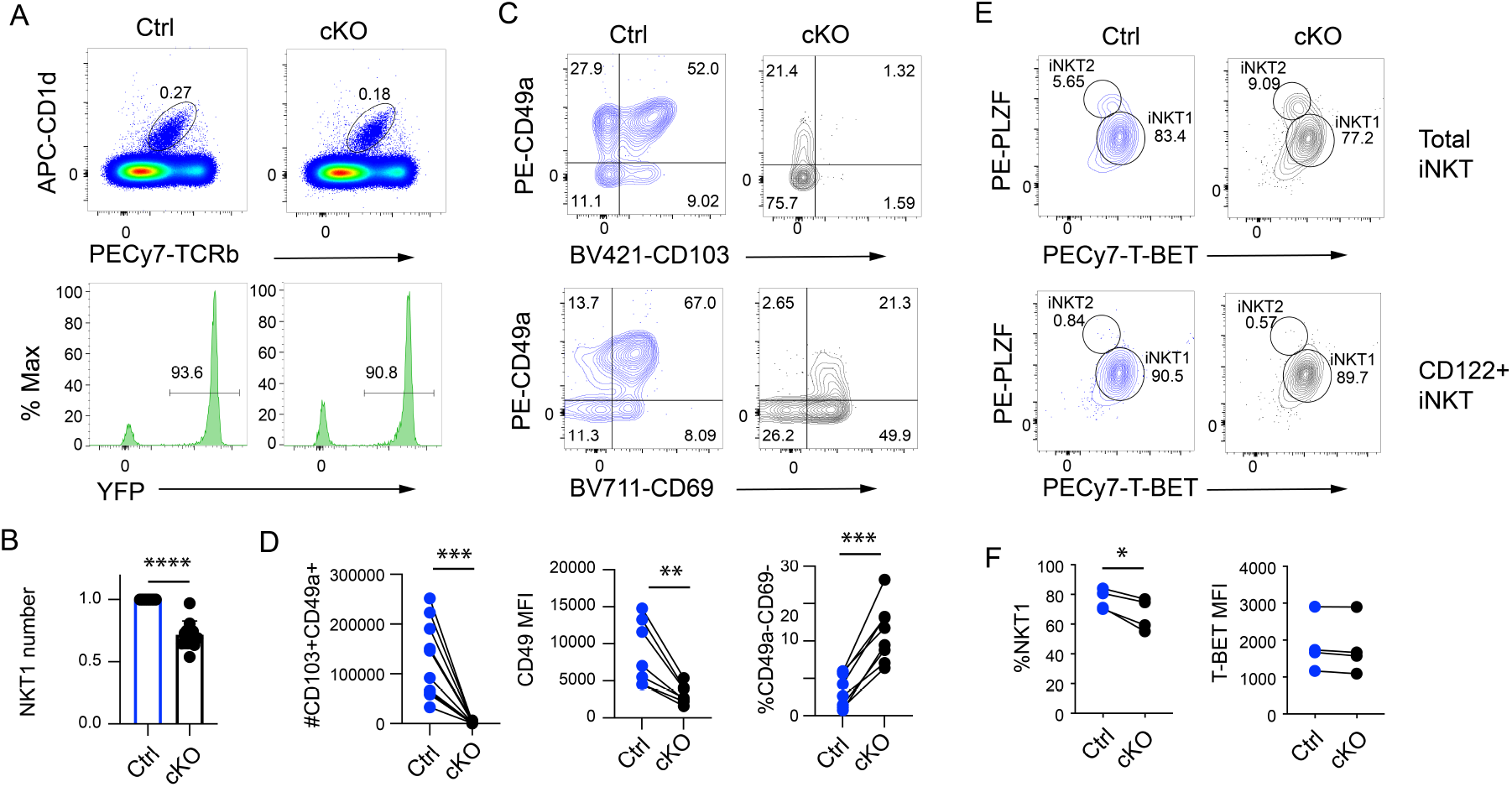
TGFβRII is required for the generation of thymic CD49a+CD103+ iNKT1 cells. (A) Flow cytometry analysis for thymic CD1d-tet^+^TCRb^+^ (top) and YFP^+^ (bottom) iNKT1 cells in Ctrl and cKO mice. (B) Summary of relative thymic iNKT1 cell numbers in Ctrl (blue) and cKO (black) mice. (C) Expression of CD49a versus CD103 (top) or CD69 (bottom) on Ctrl (blue) or cKO (black) iNKT1 cells. (D) Summary of the number of thymic CD49a+CD103+ iNKT1 cells, CD49a MFI and the percent of CD49a-CD69-iNKT1 cells. (E) Top panels: Intracellular flow cytometry for PLZF and T-BET identifying iNKT1 and iNKT2 cells among total iNKT cells in the thymus of Ctrl (blue) and cKO (black) mice. Bottom panels: PLZF and T-BET on gated CD122+ iNKT cells. (F) Summary of the % of iNKT cells that are iNKT1 among total iNKT and the MFI for T-BET in CD122^+^ iNKT cells. *P<0.05, **<P0.01, ***P<0.005. ****P<0.001 by Student’s t-test in (D) and (F) and with Welch’s test (B).

In CD8 Trm cells, CD4 Th1 cells and NK cells, TGF-β signaling dampens the expression of T-BET, despite that T-BET is required for the development of these cells (Crowl et al., 2022; Li et al., 2006b; Mackay et al., 2015; Viel et al., 2016). T-BET is also a central regulator of iNKT1 cell development (Matsuda et al., 2006; Townsend et al., 2004) and therefore we examined its expression in iNKT cells from Ctrl and cKO mice. We examined total thymic iNKT cells for expression of the signature transcription factor PLZF and T-BET, which allowed for identification of iNKT1 (T-BET^+^PLZF^lo^), iNKT2 (T-BET^-^PLZF^high^) (Kovalovsky et al., 2008; Lee et al., 2013; Savage et al., 2008). The frequency of T-BET^+^PLZF^lo^ iNKT1 cells was lower among cKO than in Ctrl iNKT cells, as expected given that there are fewer iNKT1 cells in cKO mice (Fig. 3E, F). However, regardless of whether we gated on all iNKT cells or CD122^+^ T cells, which should enrich for iNKT1 cells, the MFI of T-BET in T-BET expressing cells was not altered indicating that TGF-β was not impacting T-BET expression (Fig. 3E, F). These data demonstrate that TGFβRII is required for development of thymic iNKT1 cells and for expression of CD103 and CD49a on the surface of these cells. However, TGF-β signaling does not affect expression of T-BET in thymic iNKT cells.

### TGFβRII promotes a gene program associated with tissue residency in thymic iNKT1 cells

To gain a global view of the gene program promoted by TGF-β signaling in thymic iNKT1 cells we performed RNA-sequencing of Ctrl and *Tgfbr2*-deficient thymic iNKT1 cells. We found 110 DEGs (adj. p < 0.01), 68 that were more highly expressed in Ctrl and 42 that were more highly expressed in cKO iNKT1 cells (Fig 4A). GSEA revealed that Ctrl cells were enriched for only one pathway (FDR < 25%), the Hallmark TGF-β signaling pathway, whereas cKO cells were enriched multiple pathways including the IL-6-JAK-STAT3 pathway (Fig. 4B, S2). Genes associated with the IL6-JAK-STAT3 pathway, including *Socs3* and *Itgb3*, were both significantly higher in cKO as compared to Ctrl cells (Fig. 4C). *Tgfbr2* mRNA was confirmed to be decreased in the cKO cells as was *Smad7* mRNA, an inhibitor of TGF-β signaling induced by TGF-β (Fig. 4D). Classic targets of TGF-β signaling such as *Cdh1, Itgae, Ski, Vim*, and *Itga1* were reduced in cKO iNKT1 cells (Fig. 4E). *Inpp4b*, encoding inositol polyphospate-4-phospatase type II, which was recently implicated in TGF-β receptor endocytosis (Aki et al., 2020), was also decreased in cKO cells (Fig. S2). Multiple genes associated with migration were increased in cKO cells including *Mmp9, S1pr5, Sell* and *Hif1a* (Fig. 4F, S2). Interestingly, the gene encoding HOBIT, *Zfp683,* a transcription factor that functions redundantly with BLIMP1 to enforce tissue residency gene programs, was also decreased (Fig. 4F) (Mackay et al., 2016). However, *Prdm1* mRNA, encoding BLIMP1, was not decreased in the absence of *Tgfbr2*. *Tbx21* mRNA was not impacted, consistent with our observation that T-BET is expressed appropriately in *Tgfbr2* cKO iNKT1 cells (Fig. S2, 3E, F). *Eomes* mRNA, encoding a T box binding transcription factor related to T-BET, was very low in both Ctrl and cKO thymic iNKT1 cells (Fig. S2). Interestingly, *Zeb2*, a known T-BET target gene whose protein product regulates *S1pr5* (Dominguez et al., 2015; Evrard et al., 2022), was also not a DEG despite that *S1pr5* mRNA increased in cKO iNKT1 cells (Fig. 4F, S2). These data implicate TGFβR2 in the regulation of classic TGF-β target genes involved in adhesion and migration in iNKT1 cells and in the repression of IL6-JAK-STAT3 signaling but not in the regulation of the lineage defining transcription factor T-BET.

**Figure 4:**
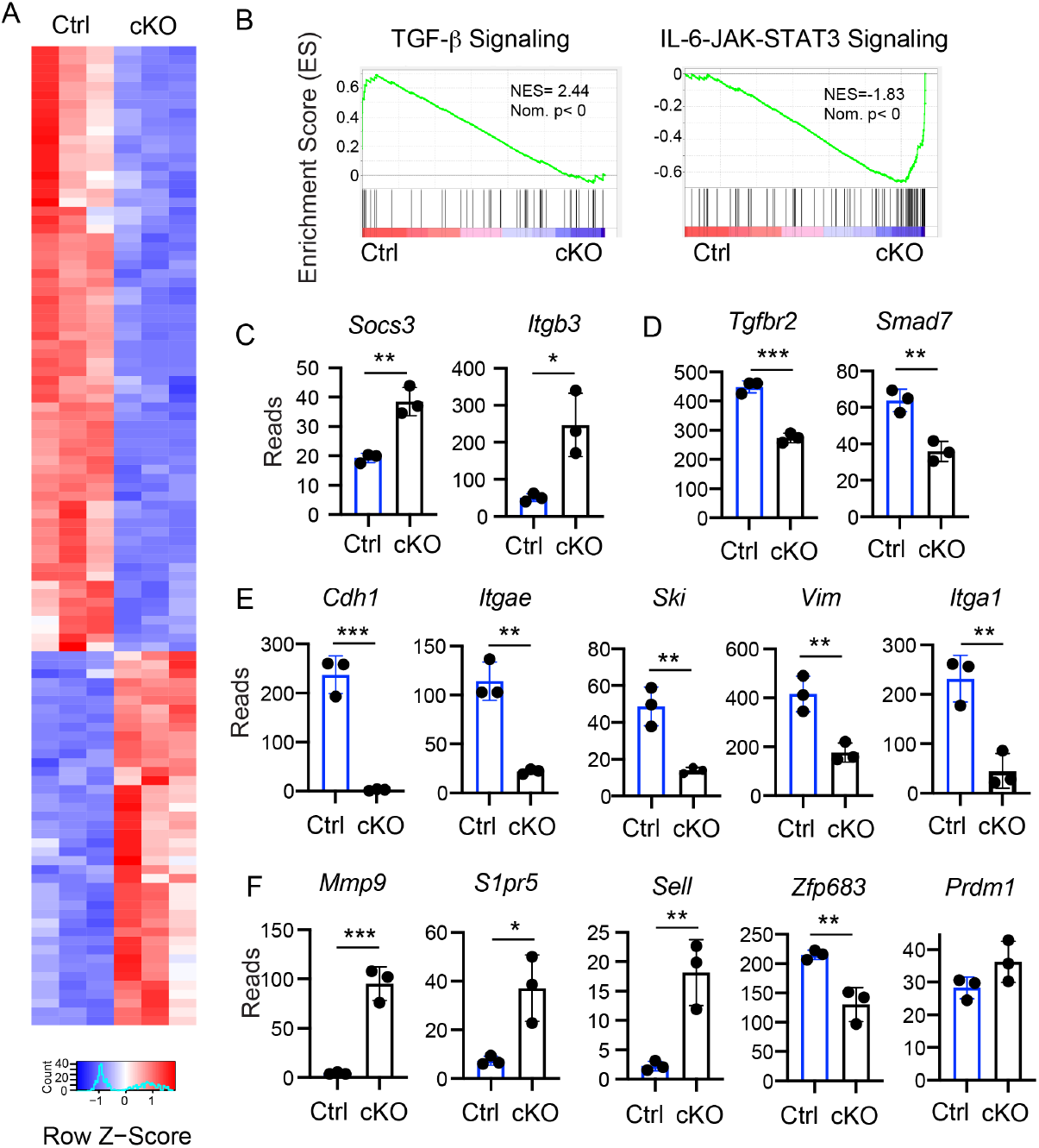
TGFβRII promotes a classic TGF-β gene signature and represses an IL6-STAT3 signaling signature in thymic iNKT1 cells. (A) Heat map for differentially expressed genes between Ctrl and cKO thymic iNKT1 by RNA-seq. (B) Top enriched Hallmark Pathway by GSEA in Ctrl and cKO. (C) Normalized reads for replicate RNA-seq samples for representative IL6-STAT3 signaling genes, (D) Tgfbr2 and Smad7, (E) classic TGF-b signaling targets and (F) genes associated with migration. *P<0.05, **<P0.01, ***P<0.005 by Student’s t-test.

The gene program regulated by TGF-β in iNKT1 cells shares significant overlap with that of skin and salivary gland CD8 Trm cells (Christo et al., 2021). However, there were also some notable differences including *Runx3, S1pr1, Junb, Cxcr4*, and *Lef1*, which were identified as TGF-β regulated in skin CD8 Trm (Christo et al., 2021) but were not observed in our dataset (Fig. S2).

### *Tgfbr2* is required for development but not thymic retention of iNKT1 cells

Thymic iNKT1 cells are tissue resident cells (Berzins et al., 2006); however, the mechanisms controlling tissue residency are not well understood. A previous study identified CD103 on thymic iNKT cells but injection of neutralizing anti-CD103 antibodies did not impact thymic retention (Wang and Hogquist, 2018). Nonetheless, we considered the hypothesis that multiple TGF-β induced genes may contribute to thymic iNKT1 residency. To test whether the decrease in thymic iNKT1 cell numbers in *Tgfbr2* cKO mice was a consequence of increased thymic emigration, we analyzed iNKT1 cell numbers after 10 days of continuous feeding of mice with the S1p lyase inhibitor 4-deoxypyridoxine (DOP) (Schwab et al., 2005). S1P lyase is essential for maintaining the gradient of S1P that promotes the emigration of thymocytes (Schwab et al., 2005). Disrupting this gradient results in retention of cells that would otherwise emigrate including mature CD4 and CD8 T cells. Treatment of Ctrl or cKO mice with DOP for 10 days resulted in an accumulation of TCRβ^+^ T cells in the thymus (Fig. S3A-C). Thymic iNKT1 cell numbers increased only in the Ctrl mice and not in cKO mice after treatment with DOP (Fig. S3 A, B, D). Total iNKT numbers did increase in the thymus of cKO mice but, in contrast to Ctrl mice, this increase was contributed almost exclusively by Rosa26-YFP negative cells (Fig. S3B). CD49a^+^CD103^+^ iNKT1 cells remained low among cKO thymic iNKT1 cells and CD49a MFI continued to be lower in cKO iNKT1 cells but CD69 continued to be expressed on the majority of cells (Fig. S3E, F). In contrast, CD49a^-^CD103^-^ (DN) iNKT1 cells accumulated in both Ctrl and cKO mice (Fig. S3G). These data are consistent with the hypothesis that CD49a^-^CD103^-^ iNKT1 cells are the most immature iNKT1 cells that either retain the potential to emigrate or are the immediate progeny of cells that had emigration potential. However, CD49a^+^ and CD49a^+^CD103^+^iNKT1 cells in Ctrl or *Tgfbr2* cKO mice do not accumulate after treatment with DOP demonstrating that iNKT1 cells are not reduced in number due to increased S1p-mediated emigration.

### *Tgfbr2* is required for optimal expression of CD49a and liver iNKT1 cell function

CD8 Trm cells in the liver do not require TGF-β but TGF-β signaling does have a subtle impact on their gene program suggesting that there is TGF-β in the liver or that these cells are imprinted prior to reaching the liver (Christo et al., 2021). Liver iNKT1 cells expressed low amounts of some TGF-β-associated genes, such as *Inpp4a* and *Vim*, and they expressed CD49a (Fig. 1E). Therefore, we examined the consequence of deleting *Tgfbr2* with *Tbx21^Cre^* on liver iNKT1 cells. In contrast to the thymus, the frequency of iNKT1 cells among liver lymphocytes was not affected in *Tgfbr2* cKO mice (Fig. 5A, B); however, the frequency of CD49a^+^ cells and the MFI of CD49a was decreased (Fig. 5C, D). CD69 continued to be expressed and the frequency of CD49a^-^CD69^-^ iNKT1 cells did not increase significantly (Fig. 5C, E), suggesting that CD49a is down regulated but the cells continue to express CD69. The observation that TGFβRII-deficiency impacted CD49a expression on liver iNKT1 cells prompted us to test the functional capacity of these cells. TGF-β impairs the functionality of CD8 Trm cells in the skin and salivary gland (Christo et al., 2021). In contrast, we found that liver iNKT1 cells from *Tgfbr2* cKO mice were somewhat less capable of producing both IFN-γ and IL-4 in response to an in vivo injection of αGalCer than liver iNKT1 cells from Ctrl mice (Fig. 5E, F). Taken together, these data demonstrate that despite the absence of a strong TGF-β gene signature in liver iNKT1 cells, TGF-β optimizes their ability to make cytokines in response to antigen stimulation.

**Figure 5:**
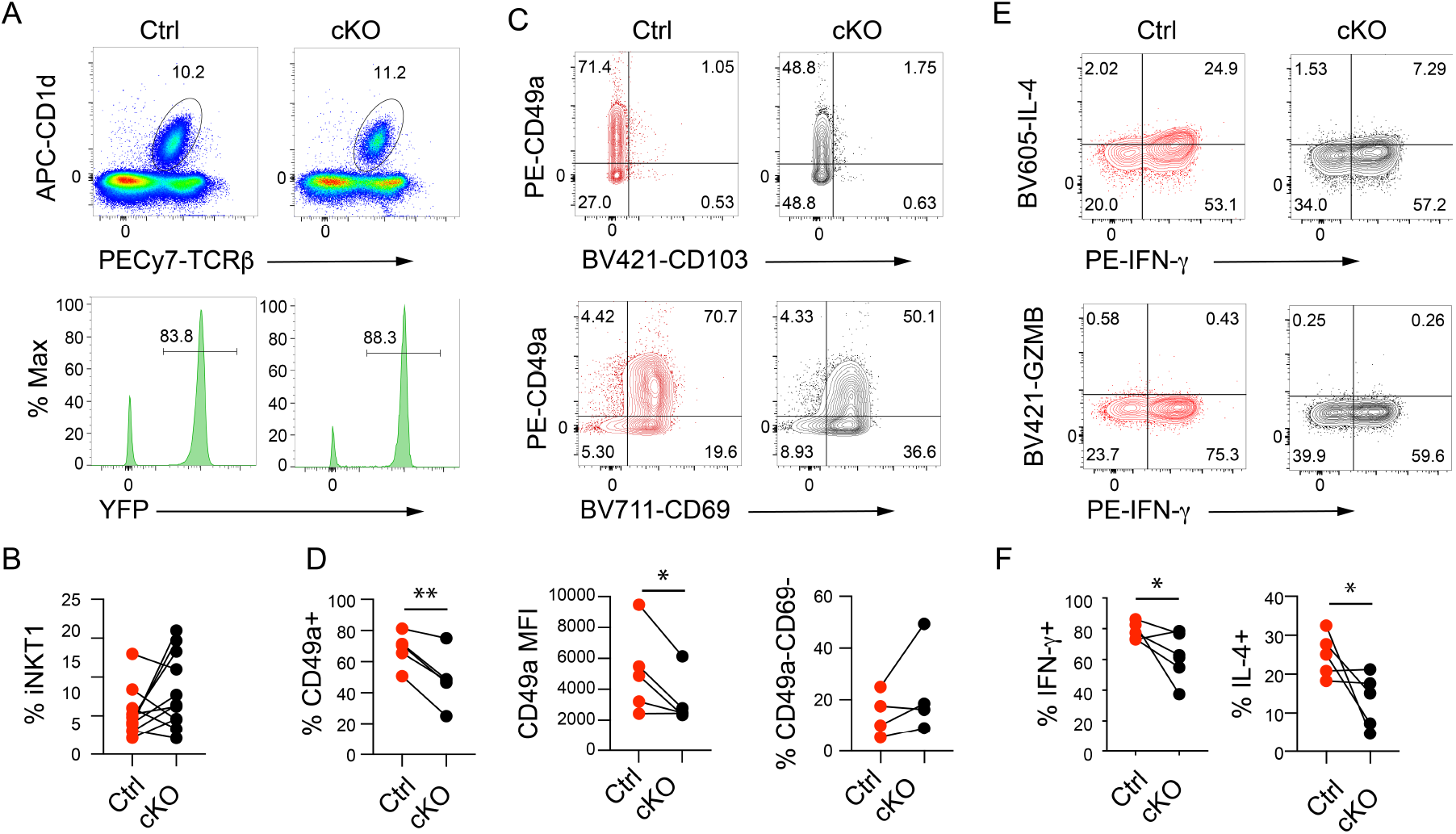
TGFβRII regulates CD49a and cytokine production from liver iNKT1 cells. (A) Flow cytometry analysis for liver CD1d-tet^+^TCRb^+^YFP^+^ iNKT1 cells in Ctrl and cKO mice. (B) Summary of the percent of iNKT1 cells among liver lymphoid cells in Ctrl (blue) and cKO (black) mice. (C) Expression of CD9a versus CD103 (top) or CD69 (bottom) on Ctrl (red) or cKO (black) liver iNKT1 cells. (D) Summary of the percent of iNKT1 cells that are CD49a^+^, the MFI of CD49, and the percent that are CD49a^-^CD69^-^. (E) Intracellular staining for IL-4 and IFN-γ (top) or GZMB and IFN-γ (bottom) and (F) summary of the percent of liver iNKT1 cells that are IFN-γ^+^ or IL-4^+^ two hours after injection of αGalCer.

These data are consistent with the possibility that TGF-β in the liver directly impacts iNKT1 cells. However, it is possible that TGF-β impacts iNKT1 cell precursors in the thymus that subsequently migrate to the liver after being imprinted by TGF-β. In this regard, we note that while iNKT1 cells that leave the thymus were proposed to be upstream of effector fate differentiation these recent thymic emigrants express detectable T-BET and could be deleted of *Tgfbr2* in our mice (Wang and Hogquist, 2018). Moreover, recent scRNA-seq studies revealed multiple iNKT cells subsets that may have the potential to emigrate from the thymus, some of which already express *Tbx21* (Harsha Krovi et al., 2020). Our DOP experiments suggest that emigrating thymic iNKT cells are CD49a^-^CD103^-^ and raise the possibility that these cells may express *Tbx21,* since some of these cells are YFP^+^ in *Tbx21^Cre^* mice. Therefore, it is feasible that emigrating iNKT1 progenitors may be imprinted by TGF-β signaling prior to leaving the thymus.

In summary, our study revealed that iNKT1 cells in the thymus differ from those in the liver and spleen by expressing genes induced by TGF-β signaling with lower expression of IL6-STAT3 signaling and fatty acid metabolism genes. We demonstrated a role for TGF-β in supporting the development of a subset of thymic iNKT1 cells that express CD49a, CD103 and CD69 while being largely dispensable for development of CD49a^+^CD103^-^ iNKT1 cells, although TGF-β does impact CD49a. The loss of TGF-β signaling did not lead to thymic iNKT1 cell emigration despite the loss of these adhesion receptors and up regulation of genes associated with emigration. However, immature thymic iNKT cells downstream of those with the potential to emigrate fail to differentiate effectively into CD49a^+^CD103^+^ cells in the absence of TGFβR2. Our data suggest that a subset of iNKT1 cells are in contact with TGF-β producing cells in the thymus, possibly medullary thymic epithelial cells (mTEC). Indeed, it is known that LTβR-dependent CCL21 ^+^ CD104^+^ MHCII^low^ mTECs impact iNKT1 cells through provision of IL-15Rα-IL-15 (Lucas et al., 2020; White et al., 2014). Moreover, Aire^+^ mTECs are dependent on CD1d, suggesting a direct role for iNKT cells in mTEC regulation, likely through provision of RANKL (White et al., 2014; White et al., 2018). Therefore, while the CD49a^+^CD103^+^subset of thymic iNKT1 cells may depend on mTECs it is also possible that they impact mTEC survival or function, a possibility that requires further exploration. Our data reveal a role for TGF-β signaling in guiding the generation of a subset of thymic iNKT1 cells and for optimal peripheral iNKT1 cell function.

## Methods

### Mice

B6.129X-Gt*(ROSA)26Sor^tm(EYFP)Cos^*/J (Rosa26-Stop-floxed-YFP), B6.CBA-Tg(Tbx21-cre)11Dlc/J (*Tbx21^Cre^*) and B6;129-*Tgfbr2^tm1Karl^*/J (*Tgfbr2^F/F^*) mice were purchased from Jackson Labs (Haddad et al., 2013; Leveen et al., 2002; Srinivas et al., 2001). All mice were housed at the University of Chicago in concordance with the guidelines of the University of Chicago Institutional Animal Care and Use Committee.

### Flow Cytometry Analysis

Mice were sacrificed using C0_2_ and thymus and liver were isolated, dissociated and passed through a mesh filter to remove debris. Liver lymphocytes were isolated using Lympholyte M (CedarLane). Cells washed twice and stained on ice for 30 minutes at a concentration of 2 × 107 cells/100 ul, washed two times and analyzed on a Fortessa flow cytometer (BD Biosciences). BV staining buffer was used when multiple BV dyes were used together (BD Biosciences). The Foxp3 staining kit was used for intracellular transcription factor staining and the Cytoperm/Cytofix Plus staining kit was used for cytokine staining (BD Biosciences). Data was analyzed using FlowJo (TreeStar). CD1d-PBS57 tetramers were titrated before use along with CD1d tetramers lacking glycolipid. Antibodies used include TCRβ, CD49a, CD103, CD69, ICAM-1, LFA-1, CD44, NK1.1, T-BET, PLZF, GZMB and IFN-γ. Fluorochromes are indicated in the flow plots.

### In vivo blocking of S1P Lyase

Mice were given DOP (30 mg/ml) in drinking water containing glucose (10 g/L) for 10 days prior to sacrifice and flow cytometry analysis (Schwab et al., 2005). Ctrl mice were given glucose containing drinking water.

### RNA-sequencing and data processing

RNA was isolated from sorted iNKT1 cells using the RNAeasy MicroKit (Qiagen). RNA libraries were constructed using Nugen’s Ovation Ultralow Library systems followed by 76 cycles of NextSeq500 sequencing (for WT thymus, liver and spleen analysis) or 100 cycles of sequencing on NovaSeq (Ctrl versus cKO analysis). Raw sequence reads were trimmed using Trimmomatic v 0.33 (TRAILING:30 MINLEN:20) and then aligned to the mouse mm10 genome with STAR v2.5.2 (Bolger et al., 2014; Trapnell et al., 2009). Reads were assigned to genes using the htseq-count tool from HTSeq v 0.6.1 and gene annotations were from Ensembl release 78 (Anders et al., 2015; Kinsella et al., 2011). The R package EdgeR was used to normalize the gene counts and to calculate differential expression statistics for each gene for each pairwise comparison of sample groups (Robinson et al., 2010). Gene set enrichment analysis was performed using gene sets from the Hallmark Pathways of MSigDB (Subramanian et al., 2005). Genes were considered differentially expressed if the fold change was > or = 2 and false discover rate <0.05. RNA-sequencing data can be accessed in the Gene Expression Omnibus (GSE215128 and GSE215338).

### Statistics

The GraphPad Prizm software package was used for statistical analysis. A Student’s t-test was used to establish the level of significance between two groups. Groups of 3 or more were assessed using ANOVA with multiple comparisons with the Geisser-Greenhouse correction. *P<0.05, **P<0.01, ***P<0.005.

## Acknowledgements

We thank E. Hegermiller and L. Lenner for technical assistance. We thank the Cytometry and Antibody Technology and Functional Genomics Core facilities at the University of Chicago. This work was supported by the National Institutes of Allergy and Infectious Diseases grant R01 AI123395 (B.L. Kee), National Cancer Institute grant P30 CA014599 (The University of Chicago Comprehensive Cancer Center) and T32 HD007009 (R.C. Morgan).

## Author Contribution

R.C. Morgan designed, performed and analyzed experiments; C.F. performed experiments; M. Sigvardsson sequenced the RNA for the wild-type comparisons; E.T. Bartom performed alignments and pipeline analysis of the RNA-seq data; B.L. Kee conceived the project, analyzed data, wrote the manuscript and obtained funding.

## Supplemental Figures

**Figure S1:**
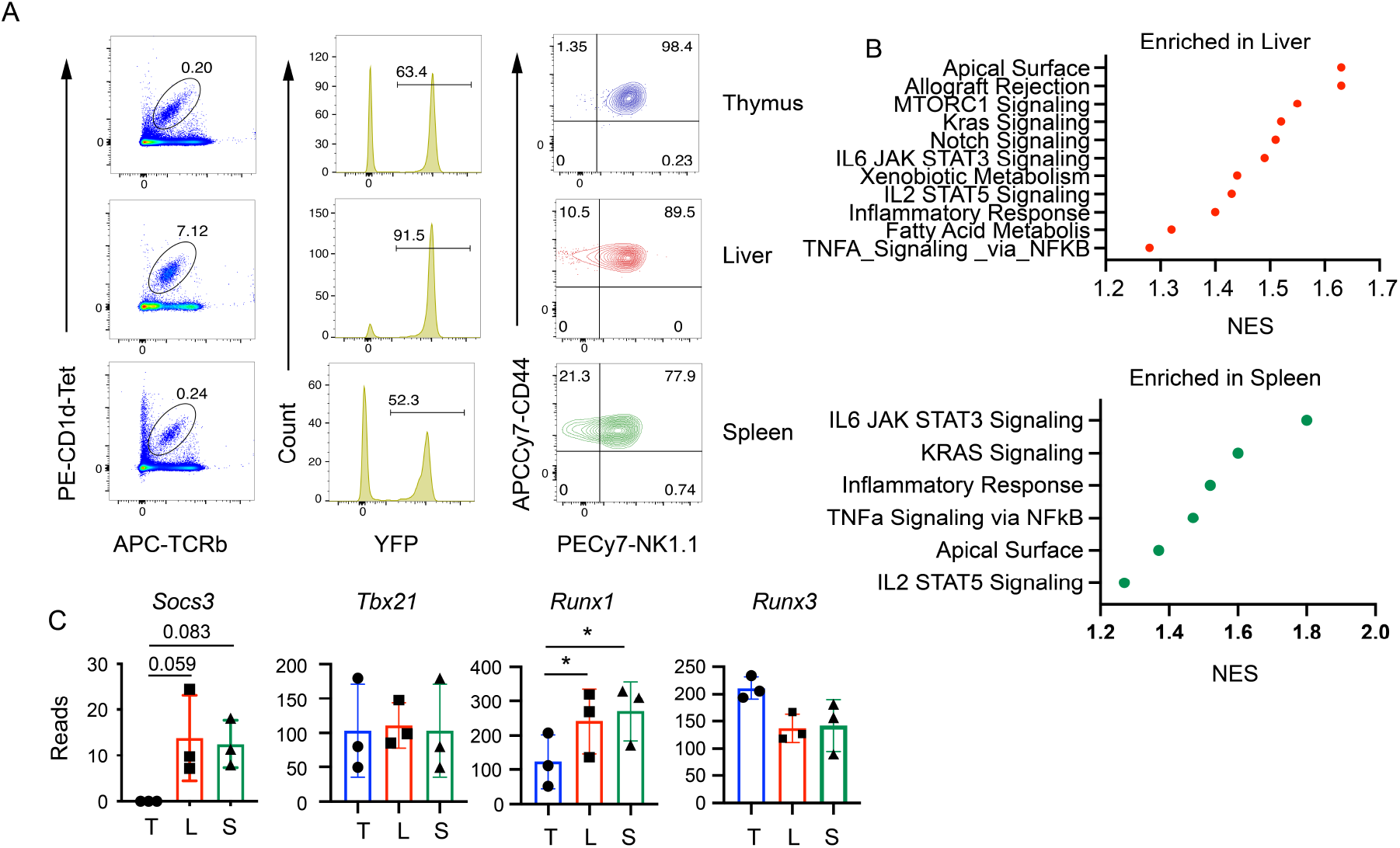
Validation of the *Tbx21^Cre^* mouse and analysis of RNA-seq data. (A) Flow cytometry analysis of NKT cells from the thymus, liver and spleen of *Tbx21^Cre^*; Rosa26-flox-stop-flox-YFP mice. iNKT cells were identified as CD1d-Tet^+^ and TCRb^+^. YFP+ iNKT cells were then examined for expression of NK1.1 and CD44. Mature iNKT1 cells express NK1.1 and progenitors that are T-BET+ can be found among CD44+NK1.1-cells. (B) Normalized Enrichment Score (NES) for genes sets with FDR <25 that were enriched in the liver (top, red) or spleen (bottom, green) compared to the thymus. (C) Normalized read counts from 3 RNA-seq samples from wild-type thymus (blue), liver (red) and spleen (green) for selected genes.

**Figure S2:**
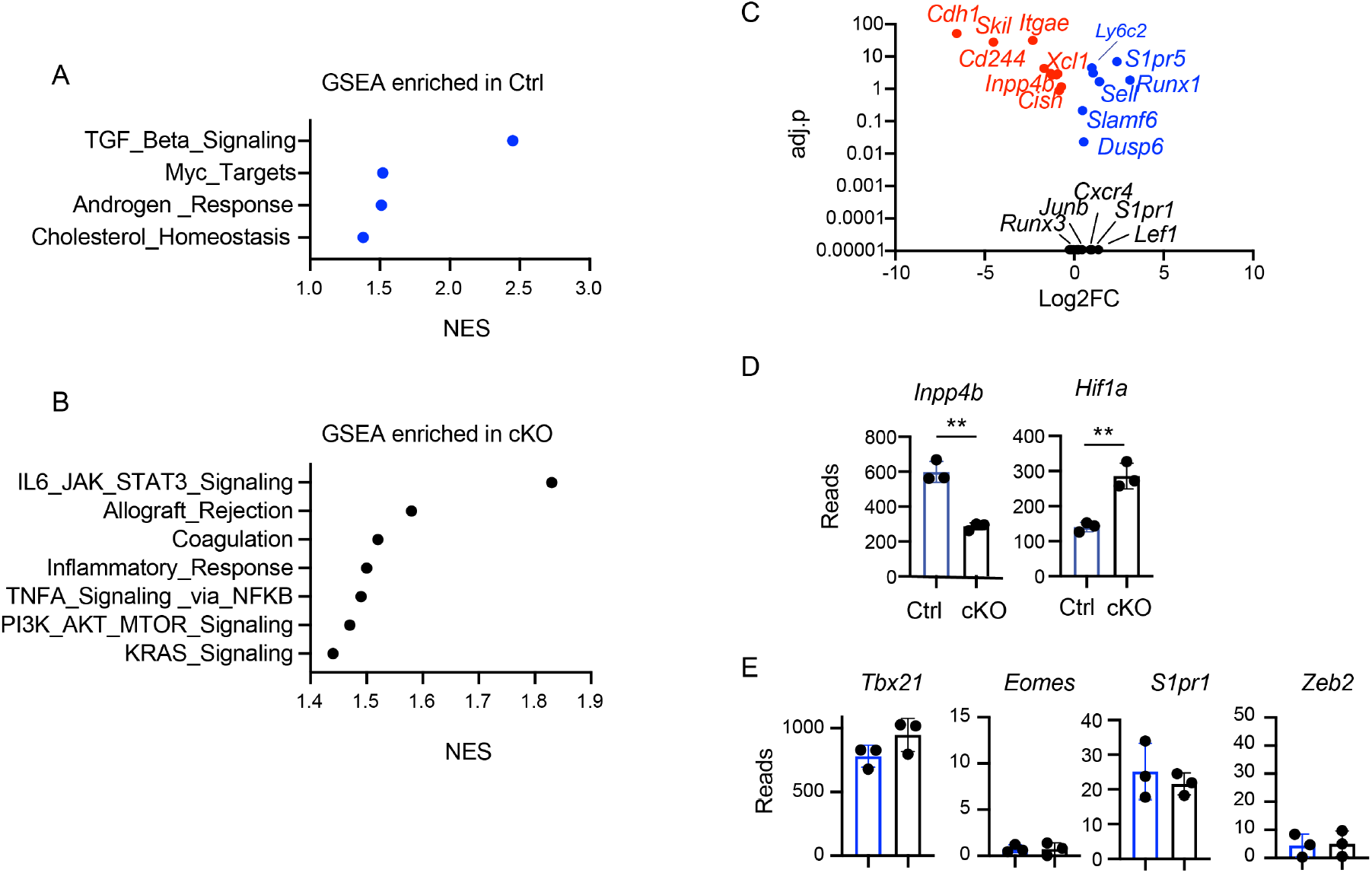
TGFβRII regulates the TGF-β signaling targets and represses cytokine associated gene expression. Normalized Enrichment Score (NES) for pathways identified by GSEA that were enriched in the thymus of (A) Ctrl or (B) *Tgfbr2* cKO iNKT1 cells. (C) Graph of Log2FC and adj.p of genes in Ctrl and cKO thymic iNKT1 cells selected for those that are TGF-β regulated in skin CD8 Trm cells (). Colors show genes that are up-regulated (blue) or down-regulated (red) or not changed (black) in our dataset. A subset of genes are labeled for clarity. (D) Normalized read counts for *Inpp4b* and *Hif1a*, or (E) *Tbx21, Eomes, S1pr1* and *Zeb2* in Ctrl (blue) and cKO (black) thymic iNKT1 cells by RNA-seq. Each circle is one RNA-seq replicate.

**Figure S3:**
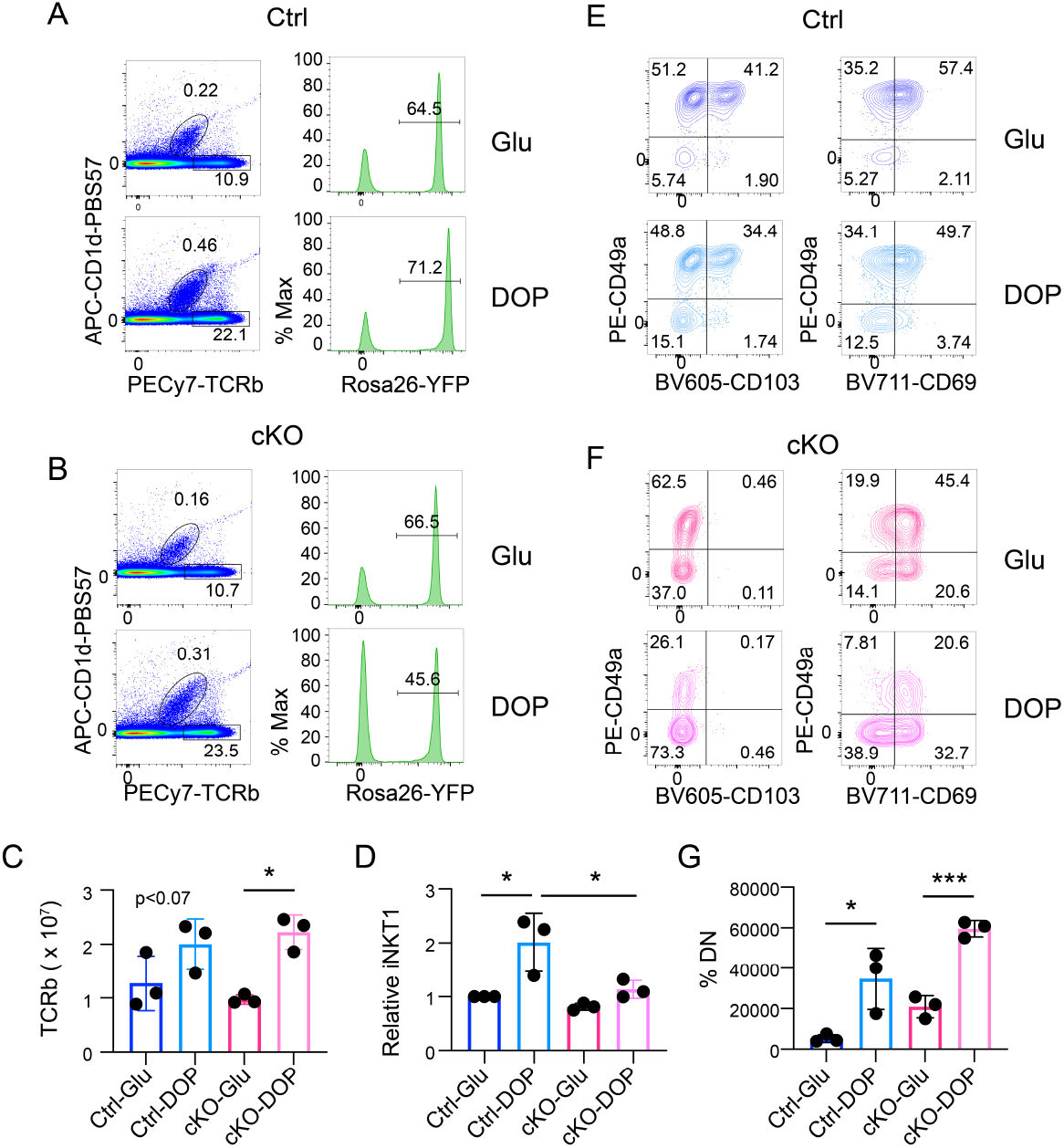
Inhibition of the S1P-gradient reveals that TGF-β does support iNKT1 cells numbers by preventing thymic emigration. Flow cytometry was performed to identify TCRb^+^ T cells or CD1d-tet^+^TCRb^+^YFP^+^ iNKT1 cells in Ctrl (A) and cKO (B) mice fed glucose (Glu) or DPO (DOP) for 10 days. (C) Summary of the number of TCRb+ T cells or the (D) relative number of iNKT1 cells in Ctrl (blue) or cKO (pink) mice fed Glu or DOP. (E) FACS plots showing expression of CD49a versus CD103 or CD69 on Glu (upper) or DOP (lower) fed Ctrl or (F) cKO mice. (G) The percent of iNKT1 cells that are negative for CD49a and CD69 (DN) in Ctrl or cKO mice fed Glu or DOP. *P<0.05, **<P0.01, ***P<0.005 by ANOVA with multiple comparisons.

## References

Aki, S., K. Yoshioka, N. Takuwa, and Y. Takuwa. 2020. TGFbeta receptor endocytosis and Smad signaling require synaptojanin1, PI3K-C2alpha-, and INPP4B-mediated phosphoinositide conversions. Mol Biol Cell 31:360–372.

Anders, S., P.T. Pyl, and W. Huber. 2015. HTSeq--a Python framework to work with high-throughput sequencing data. Bioinformatics 31:166–169.

Baranek, T., K. Lebrigand, C. de Amat Herbozo, L. Gonzalez, G. Bogard, C. Dietrich, V. Magnone, C. Boisseau, Y. Jouan, F. Trottein, M. Si-Tahar, M. Leite-de-Moraes, T. Mallevaey, and C. Paget. 2020. High Dimensional Single-Cell Analysis Reveals iNKT Cell Developmental Trajectories and Effector Fate Decision. Cell Rep 32:108116.

Bendelac, A., P.B. Savage, and L. Teyton. 2007. The biology of NKT cells. Annu Rev Immunol 25:297–336.

Benlagha, K., T. Kyin, A. Beavis, L. Teyton, and A. Bendelac. 2002. A thymic precursor to the NK T cell lineage. Science 296:553–555.

Benlagha, K., D.G. Wei, J. Veiga, L. Teyton, and A. Bendelac. 2005. Characterization of the early stages of thymic NKT cell development. J Exp Med 202:485–492.

Berzins, S.P., F.W. McNab, C.M. Jones, M.J. Smyth, and D.I. Godfrey. 2006. Long-term retention of mature NK1.1+ NKT cells in the thymus. J Immunol 176:4059–4065.

Bolger, A.M., M. Lohse, and B. Usadel. 2014. Trimmomatic: a flexible trimmer for Illumina sequence data. Bioinformatics 30:2114–2120.

Breed, E.R., M. Voboril, K.M. Ashby, R.J. Martinez, L. Qian, H. Wang, O.C. Salgado, C.H. O’Connor, and K.A. Hogquist. 2022. Type 2 cytokines in the thymus activate Sirpalpha(+) dendritic cells to promote clonal deletion. Nat Immunol 23:1042–1051.

Christo, S.N., M. Evrard, S.L. Park, L.C. Gandolfo, T.N. Burn, R. Fonseca, D.M. Newman, Y.O. Alexandre, N. Collins, N.M. Zamudio, F. Souza-Fonseca-Guimaraes, D.G. Pellicci, D. Chisanga, W. Shi, L. Bartholin, G.T. Belz, N.D. Huntington, A. Lucas, M. Lucas, S.N. Mueller, W.R. Heath, F. Ginhoux, T.P. Speed, F.R. Carbone, A. Kallies, and L.K. Mackay. 2021. Discrete tissue microenvironments instruct diversity in resident memory T cell function and plasticity. Nat Immunol 22:1140–1151.

Crosby, C.M., and M. Kronenberg. 2018. Tissue-specific functions of invariant natural killer T cells. Nat Rev Immunol 18:559–574.

Crowl, J.T., M. Heeg, A. Ferry, J.J. Milner, K.D. Omilusik, C. Toma, Z. He, J.T. Chang, and A.W. Goldrath. 2022. Tissue-resident memory CD8(+) T cells possess unique transcriptional, epigenetic and functional adaptations to different tissue environments. Nat Immunol 23:1121–1131.

Doisne, J.M., L. Bartholin, K.P. Yan, C.N. Garcia, N. Duarte, J.B. Le Luduec, D. Vincent, F. Cyprian, B. Horvat, S. Martel, R. Rimokh, R. Losson, K. Benlagha, and J.C. Marie. 2009. iNKT cell development is orchestrated by different branches of TGF-beta signaling. J Exp Med 206:1365–1378.

Dominguez, C.X., R.A. Amezquita, T. Guan, H.D. Marshall, N.S. Joshi, S.H. Kleinstein, and S.M. Kaech. 2015. The transcription factors ZEB2 and T-bet cooperate to program cytotoxic T cell terminal differentiation in response to LCMV viral infection. J Exp Med 212:2041–2056.

Engel, I., G. Seumois, L. Chavez, D. Samaniego-Castruita, B. White, A. Chawla, D. Mock, P. Vijayanand, and M. Kronenberg. 2016. Innate-like functions of natural killer T cell subsets result from highly divergent gene programs. Nat Immunol 17:728–739.

Evrard, M., E. Wynne-Jones, C. Peng, Y. Kato, S.N. Christo, R. Fonseca, S.L. Park, T.N. Burn, M. Osman, S. Devi, J. Chun, S.N. Mueller, G. Kannourakis, S.P. Berzins, D.G. Pellicci, W.R. Heath, S.C. Jameson, and L.K. Mackay. 2022. Sphingosine 1-phosphate receptor 5 (S1PR5) regulates the peripheral retention of tissue-resident lymphocytes. J Exp Med 219:

Fonseca, R., T.N. Burn, L.C. Gandolfo, S. Devi, S.L. Park, A. Obers, M. Evrard, S.N. Christo, F.A. Buquicchio, C.A. Lareau, K.M. McDonald, S.K. Sandford, N.M. Zamudio, N.G. Zanluqui, A. Zaid, T.P. Speed, A.T. Satpathy, S.N. Mueller, F.R. Carbone, and L.K. Mackay. 2022. Runx3 drives a CD8(+) T cell tissue residency program that is absent in Cd4(+) T cells. Nat Immunol 23:1236–1245.

Frizzell, H., R. Fonseca, S.N. Christo, M. Evrard, S. Cruz-Gomez, N.G. Zanluqui, B. von Scheidt, D. Freestone, S.L. Park, H.E.G. McWilliam, J.A. Villadangos, F.R. Carbone, and L.K. Mackay. 2020. Organ-specific isoform selection of fatty acid-binding proteins in tissue-resident lymphocytes. Sci Immunol 5:

Haddad, R., A. Lanjuin, L. Madisen, H. Zeng, V.N. Murthy, and N. Uchida. 2013. Olfactory cortical neurons read out a relative time code in the olfactory bulb. Nat Neurosci 16:949–957.

Hadley, G.A., E.A. Rostapshova, D.M. Gomolka, B.M. Taylor, S.T. Bartlett, C.I. Drachenberg, and M.R. Weir. 1999. Regulation of the epithelial cell-specific integrin, CD103, by human CD8+ cytolytic T lymphocytes. Transplantation 67:1418–1425.

Harsha Krovi, S., J. Zhang, M.J. Michaels-Foster, T. Brunetti, L. Loh, J. Scott-Browne, and L. Gapin. 2020. Thymic iNKT single cell analyses unmask the common developmental program of mouse innate T cells. Nat Commun 11:6238.

Havenar-Daughton, C., S. Li, K. Benlagha, and J.C. Marie. 2012. Development and function of murine RORgammat+ iNKT cells are under TGF-beta signaling control. Blood 119:3486–3494.

Kinsella, R.J., A. Kahari, S. Haider, J. Zamora, G. Proctor, G. Spudich, J. Almeida-King, D. Staines, P. Derwent, A. Kerhornou, P. Kersey, and P. Flicek. 2011. Ensembl BioMarts: a hub for data retrieval across taxonomic space. Database (Oxford) 2011:bar030.

Kovalovsky, D., O.U. Uche, S. Eladad, R.M. Hobbs, W. Yi, E. Alonzo, K. Chua, M. Eidson, H.J. Kim, J.S. Im, P.P. Pandolfi, and D.B. Sant’Angelo. 2008. The BTB-zinc finger transcriptional regulator PLZF controls the development of invariant natural killer T cell effector functions. Nat Immunol 9:1055–1064.

Lee, Y.J., K.L. Holzapfel, J. Zhu, S.C. Jameson, and K.A. Hogquist. 2013. Steady-state production of IL-4 modulates immunity in mouse strains and is determined by lineage diversity of iNKT cells. Nat Immunol 14:1146–1154.

Lee, Y.J., G.J. Starrett, S.T. Lee, R. Yang, C.M. Henzler, S.C. Jameson, and K.A. Hogquist. 2016. Lineage-Specific Effector Signatures of Invariant NKT Cells Are Shared amongst gammadelta T, Innate Lymphoid, and Th Cells. J Immunol 197:1460–1470.

Lee, Y.J., H. Wang, G.J. Starrett, V. Phuong, S.C. Jameson, and K.A. Hogquist. 2015. Tissue-Specific Distribution of iNKT Cells Impacts Their Cytokine Response. Immunity 43:566–578.

Leveen, P., J. Larsson, M. Ehinger, C.M. Cilio, M. Sundler, L.J. Sjostrand, R. Holmdahl, and S. Karlsson. 2002. Induced disruption of the transforming growth factor beta type II receptor gene in mice causes a lethal inflammatory disorder that is transplantable. Blood 100:560–568.

Li, M.O., S. Sanjabi, and R.A. Flavell. 2006a. Transforming growth factor-beta controls development, homeostasis, and tolerance of T cells by regulatory T cell-dependent and -independent mechanisms. Immunity 25:455–471.

Li, M.O., Y.Y. Wan, S. Sanjabi, A.K. Robertson, and R.A. Flavell. 2006b. Transforming growth factor-beta regulation of immune responses. Annu Rev Immunol 24:99–146.

Lucas, B., A.J. White, E.J. Cosway, S.M. Parnell, K.D. James, N.D. Jones, I. Ohigashi, Y. Takahama, W.E. Jenkinson, and G. Anderson. 2020. Diversity in medullary thymic epithelial cells controls the activity and availability of iNKT cells. Nat Commun 11:2198.

Mackay, L.K., M. Minnich, N.A. Kragten, Y. Liao, B. Nota, C. Seillet, A. Zaid, K. Man, S. Preston, D. Freestone, A. Braun, E. Wynne-Jones, F.M. Behr, R. Stark, D.G. Pellicci, D.I. Godfrey, G.T. Belz, M. Pellegrini, T. Gebhardt, M. Busslinger, W. Shi, F.R. Carbone, R.A. van Lier, A. Kallies, and K.P. van Gisbergen. 2016. Hobit and Blimp1 instruct a universal transcriptional program of tissue residency in lymphocytes. Science 352:459–463.

Mackay, L.K., A. Rahimpour, J.Z. Ma, N. Collins, A.T. Stock, M.L. Hafon, J. Vega-Ramos, P. Lauzurica, S.N. Mueller, T. Stefanovic, D.C. Tscharke, W.R. Heath, M. Inouye, F.R. Carbone, and T. Gebhardt. 2013. The developmental pathway for CD103(+)CD8+ tissue-resident memory T cells of skin. Nat Immunol 14:1294–1301.

Mackay, L.K., E. Wynne-Jones, D. Freestone, D.G. Pellicci, L.A. Mielke, D.M. Newman, A. Braun, F. Masson, A. Kallies, G.T. Belz, and F.R. Carbone. 2015. T-box Transcription Factors Combine with the Cytokines TGF-beta and IL-15 to Control Tissue-Resident Memory T Cell Fate. Immunity 43:1101–1111.

Matsuda, J.L., Q. Zhang, R. Ndonye, S.K. Richardson, A.R. Howell, and L. Gapin. 2006. T-bet concomitantly controls migration, survival, and effector functions during the development of Valpha14i NKT cells. Blood 107:2797–2805.

Mokrani, M., J. Klibi, D. Bluteau, G. Bismuth, and F. Mami-Chouaib. 2014. Smad and NFAT pathways cooperate to induce CD103 expression in human CD8 T lymphocytes. J Immunol 192:2471–2479.

Murray, M.P., I. Engel, G. Seumois, S. Herrera-De la Mata, S.L. Rosales, A. Sethi, A. Logandha Ramamoorthy Premlal, G.Y. Seo, J. Greenbaum, P. Vijayanand, J.P. Scott-Browne, and M. Kronenberg. 2021. Transcriptome and chromatin landscape of iNKT cells are shaped by subset differentiation and antigen exposure. Nat Commun 12:1446.

Nath, A.P., A. Braun, S.C. Ritchie, F.R. Carbone, L.K. Mackay, T. Gebhardt, and M. Inouye. 2019. Comparative analysis reveals a role for TGF-beta in shaping the residency-related transcriptional signature in tissue-resident memory CD8+ T cells. PLoS One 14:e0210495.

Pan, Y., T. Tian, C.O. Park, S.Y. Lofftus, S. Mei, X. Liu, C. Luo, J.T. O’Malley, A. Gehad, J.E. Teague, S.J. Divito, R. Fuhlbrigge, P. Puigserver, J.G. Krueger, G.S. Hotamisligil, R.A. Clark, and T.S. Kupper. 2017. Survival of tissue-resident memory T cells requires exogenous lipid uptake and metabolism. Nature 543:252–256.

Robinson, M.D., D.J. McCarthy, and G.K. Smyth. 2010. edgeR: a Bioconductor package for differential expression analysis of digital gene expression data. Bioinformatics 26:139–140.

Savage, A.K., M.G. Constantinides, J. Han, D. Picard, E. Martin, B. Li, O. Lantz, and A. Bendelac. 2008. The transcription factor PLZF directs the effector program of the NKT cell lineage. Immunity 29:391–403.

Schwab, S.R., J.P. Pereira, M. Matloubian, Y. Xu, Y. Huang, and J.G. Cyster. 2005. Lymphocyte sequestration through S1P lyase inhibition and disruption of S1P gradients. Science 309:1735–1739.

Shiow, L.R., D.B. Rosen, N. Brdickova, Y. Xu, J. An, L.L. Lanier, J.G. Cyster, and M. Matloubian. 2006. CD69 acts downstream of interferon-alpha/beta to inhibit S1P1 and lymphocyte egress from lymphoid organs. Nature 440:540–544.

Shissler, S.C., and T.J. Webb. 2019. The ins and outs of type I iNKT cell development. Mol Immunol 105:116–130.

Srinivas, S., T. Watanabe, C.S. Lin, C.M. William, Y. Tanabe, T.M. Jessell, and F. Costantini. 2001. Cre reporter strains produced by targeted insertion of EYFP and ECFP into the ROSA26 locus. BMC Dev Biol 1:4.

Stein, J.V., N. Ruef, and S. Wissmann. 2021. Organ-Specific Surveillance and Long-Term Residency Strategies Adapted by Tissue-Resident Memory CD8(+) T Cells. Front Immunol 12:626019.

Subramanian, A., P. Tamayo, V.K. Mootha, S. Mukherjee, B.L. Ebert, M.A. Gillette, A. Paulovich, S.L. Pomeroy, T.R. Golub, E.S. Lander, and J.P. Mesirov. 2005. Gene set enrichment analysis: a knowledge-based approach for interpreting genomewide expression profiles. Proc Natl Acad Sci U S A 102:15545–15550.

Syn, W.K., K.M. Agboola, M. Swiderska, G.A. Michelotti, E. Liaskou, H. Pang, G. Xie, G. Philips, I.S. Chan, G.F. Karaca, A. Pereira Tde, Y. Chen, Z. Mi, P.C. Kuo, S.S. Choi, C.D. Guy, M.F. Abdelmalek, and A.M. Diehl. 2012. NKT-associated hedgehog and osteopontin drive fibrogenesis in non-alcoholic fatty liver disease. Gut 61:1323–1329.

Syn, W.K., Y.H. Oo, T.A. Pereira, G.F. Karaca, Y. Jung, A. Omenetti, R.P. Witek, S.S. Choi, C.D. Guy, C.M. Fearing, V. Teaberry, F.E. Pereira, D.H. Adams, and A.M. Diehl. 2010. Accumulation of natural killer T cells in progressive nonalcoholic fatty liver disease. Hepatology 51:1998–2007.

Thomas, S.Y., S.T. Scanlon, K.G. Griewank, M.G. Constantinides, A.K. Savage, K.A. Barr, F. Meng, A.D. Luster, and A. Bendelac. 2011. PLZF induces an intravascular surveillance program mediated by long-lived LFA-1-ICAM-1 interactions. J Exp Med 208:1179–1188.

Townsend, M.J., A.S. Weinmann, J.L. Matsuda, R. Salomon, P.J. Farnham, C.A. Biron, L. Gapin, and L.H. Glimcher. 2004. T-bet regulates the terminal maturation and homeostasis of NK and Valpha14i NKT cells. Immunity 20:477–494.

Trapnell, C., L. Pachter, and S.L. Salzberg. 2009. TopHat: discovering splice junctions with RNA-Seq. Bioinformatics 25:1105–1111.

Umeshappa, C.S., P. Sole, J. Yamanouchi, S. Mohapatra, B.G.J. Surewaard, J. Garnica, S. Singha, D. Mondal, E. Cortes-Vicente, C. D’Mello, A. Mason, P. Kubes, P. Serra, Y. Yang, and P. Santamaria. 2022. Re-programming mouse liver-resident invariant natural killer T cells for suppressing hepatic and diabetogenic autoimmunity. Nat Commun 13:3279.

Verykokakis, M., M.D. Boos, A. Bendelac, and B.L. Kee. 2010. SAP protein-dependent natural killer T-like cells regulate the development of CD8(+) T cells with innate lymphocyte characteristics. Immunity 33:203–215.

Verykokakis, M., E.C. Zook, and B.L. Kee. 2014. ID’ing innate and innate-like lymphoid cells. Immunol Rev 261:177–197.

Viel, S., A. Marcais, F.S. Guimaraes, R. Loftus, J. Rabilloud, M. Grau, S. Degouve, S. Djebali, A. Sanlaville, E. Charrier, J. Bienvenu, J.C. Marie, C. Caux, J. Marvel, L. Town, N.D. Huntington, L. Bartholin, D. Finlay, M.J. Smyth, and T. Walzer. 2016. TGF-beta inhibits the activation and functions of NK cells by repressing the mTOR pathway. Sci Signal 9:ra19.

Wang, H., and K.A. Hogquist. 2018. CCR7 defines a precursor for murine iNKT cells in thymus and periphery. Elife 7:

Weinreich, M.A., O.A. Odumade, S.C. Jameson, and K.A. Hogquist. 2010. T cells expressing the transcription factor PLZF regulate the development of memorylike CD8+ T cells. Nat Immunol 11:709–716.

White, A.J., W.E. Jenkinson, J.E. Cowan, S.M. Parnell, A. Bacon, N.D. Jones, E.J. Jenkinson, and G. Anderson. 2014. An essential role for medullary thymic epithelial cells during the intrathymic development of invariant NKT cells. J Immunol 192:2659–2666.

White, A.J., B. Lucas, W.E. Jenkinson, and G. Anderson. 2018. Invariant NKT Cells and Control of the Thymus Medulla. J Immunol 200:3333–3339.

Zhang, N., and M.J. Bevan. 2013. Transforming growth factor-beta signaling controls the formation and maintenance of gut-resident memory T cells by regulating migration and retention. Immunity 39:687–696.

